# Trait Correlates of Climatic Niche Tracking in British Birds

**DOI:** 10.1101/103507

**Authors:** Giovanni Rapacciuolo, Robert A. Robinson, Simon Gillings, Andy Purvis

## Abstract

Growing evidence indicates that species respond idiosyncratically when exposed to the same changes in climate. As a result, understanding the potential influence of biological traits on species’ distributional responses is a research priority. Yet, empirical support for hypothesised influences of traits on climate change responses remains equivocal.

In this paper, we developed a novel approach to determine whether biological traits predict the degree of climatic niche tracking of British breeding birds in response to recent climate change. First, we quantified how well predicted positive and negative changes in probability of presence from climate-based species distribution models agreed with observed local gains and losses in species’ occupancy – our measure of climatic niche tracking. Second, we examined whether the degree of climatic niche tracking could be predicted by species’ ecological and life-history traits, as well as phylogenetic relationships.

Overall, British breeding birds displayed a low degree of climatic niche tracking over the period of our study, though this varied substantially among species. Models incorporating traits and phylogeny explained a low proportion of the variation in climatic niche tracking. Nevertheless, we did find statistical evidence that species with lower lifespans tracked their climatic niches more closely, whilst species with a mixed diet displayed a lower degree of climatic niche tracking.

We present here a tractable approach for quantifying the degree to which observed local range gains and losses can be related to climate redistribution and apply it to British breeding birds. Although we do not find strong evidence that traits predict the degree of climatic niche tracking, we discuss why this is likely to be a consequence of the features of our study system rather than the approach itself. We believe this approach may prove to be useful as datasets of temporal changes in species distributions become increasingly available.

## INTRODUCTION

Global climatic conditions are changing rapidly and further dramatic changes are projected for this century (IPCC, 2013). Spatial and temporal variability in rates of change lead to the continuous redistribution of climatic conditions across the globe (Loarie *et al*., 2009). If species have evolved physiological adaptations to local climatic conditions (Phillimore *et al*., 2010), they may respond to climate change by either migrating to track their existing climatic associations, persisting *in situ* within altered climatic conditions through plasticity or adaptation, or becoming locally extinct (La Sorte & Jetz, 2012). Understanding how species respond to climate redistribution is critical for improving our forecasts of species’ future responses and the conservation value of our mitigation actions.

It is now clear that animal species are responding idiosyncratically to changes in climate (Chen *et al*., 2011; Rapacciuolo *et al*., 2014a), as they did in the Pleistocene (Stewart, 2008; Hofreiter & Stewart, 2009). As a result, a growing body of theory focuses on the potential influence of biological traits on the distributional responses of species to climate change (Williams *et al*., 2008; Foden *et al*., 2013). While substantial progress in this area has been achieved for ectothermic vertebrates (Buckley, 2010; Huey *et al*., 2012), empirical support toward trait-based hypotheses of climate change responses in endotherms remains contrasting (Angert *et al*., 2011; Cahill *et al*., 2013; McCain & King, 2014). One reason for this may be that the majority of studies to date have focused on related but slightly different questions. Some have focused on trait correlates of overall distributional response or vulnerability, without specific attention to climate responses (Angert *et al*., 2011; Pocock, 2011; Bradshaw *et al*., 2014). These studies are of limited use for understanding trait effects on climate responses since these responses are confounded with responses to additional synergistic drivers of change. Other studies, while focusing on climatic associations, typically focus on changes in species’ geographic ranges as a whole (Kharouba *et al*. 2009, Dobrowski *et al*. 2011, Smith *et al*. 2013; but see McCain and King 2014). However, migration, persistence, and extinction are not mutually exclusive responses to climate change across the range of a single species (Tingley *et al*., 2012; Rapacciuolo *et al*., 2014a; Rowe *et al*., 2014). Instead, overall species’ trends result from the net demographic impacts of these three possible responses (Angert *et al*., 2011). Identifying local responses independent of overall trends is a crucial step towards a comprehensive spatially-explicit assessment of species’ vulnerability to climate change. This is especially important given that synergistic drivers of change (e.g. land use change and extreme disturbance events) also impact biodiversity heterogeneously across space and may exacerbate local vulnerability (Turner, 2010).

In this paper, we overcome some of the limitations of existing studies by using a recently-published method (Rapacciuolo *et al*., 2014b) to spatially quantify the agreement between observed range changes and predictions based on climate redistribution – a spatially-explicit measure of climate niche tracking. Our objectives were to examine whether British breeding birds are tracking their climatic niches over time and whether biological traits are related to the degree of climatic niche tracking. First, we built climate-based species distribution models and generated predictions of change in the probability of presence of bird species based on the redistribution of each species’ historical climatic niche across Great Britain. Second, we quantified how the agreement between these predictions and observed species’ gains and losses over the same time interval varied spatially throughout species’ geographic ranges. British breeding birds are one of only a handful of systems enabling such analyses at a large spatial scale, since their distributions have been sampled comprehensively at repeated time intervals across all of Great Britain’s 10-km Ordnance Survey National Grid squares (Sharrock, 1976; Gibbons *et al*., 1993). Given this unusually-constant sampling effort over time and space, we were able to derive estimates of local range gains and losses over an approximately 30-year period and relate them to climate redistribution over the same spatial and temporal scale.

Finally, because the degree to which species track their climatic conditions is likely to depend on their particular ecological and life-history traits (Williams *et al*., 2008; Huey *et al*., 2012; Foden *et al*., 2013), we tested four hypotheses of the effect of traits on climatic niche tracking. We hypothesised that: (i) more mobile species which can disperse greater distances would be better able to track their climatic niches across newly-suitable areas (Schloss *et al*., 2012); (ii) species with faster life histories would be better able to track their climatic niches due to their higher intrinsic rate of population growth and resulting ability to recover quickly from low numbers (Angert *et al*., 2011; Auer & King, 2014); (iii) habitat specialists would be less able to track their climatic niches given their greater difficulty in establishing populations in new habitats and/or keeping high numbers under altered conditions (Angert *et al*., 2011). (iv) higher-trophic-level species would display less climatic niche tracking given the higher number of trophic links separating them from the direct effects of climate on primary producers (Huntley *et al*., 2004). Furthermore, we tested for phylogenetic signal in climatic niche tracking in order to assess whether additional attributes of species not captured by our traits could be associated with variation in climatic niche tracking.

## METHODS

### Species distribution data

We used occupancy records for 226 British breeding birds at a 10-km grid square resolution in two time periods of intensive recording effort (*t*_*1*_: 1968–1972; *t*_*2*_: 1988–1991), each leading to the publication of a national breeding bird atlas (Sharrock, 1976; Gibbons *et al*., 1993). To avoid problems related to building models with extremely small sample sizes (Wisz *et al*., 2008), we excluded 43 species occupying fewer than 20 grid squares in either time period. We excluded a further 71 predominantly-aquatic species (i.e. marine birds, waterfowl, and shorebirds), given the substantial difficulties in defining local range gains/losses for these species. Although species’ absence from each 10–km grid square was not recorded during sampling, 98 – 100% grid squares in Great Britain were sampled meticulously during both time periods, with high levels of replicate recording and under-recorded areas targeted by extra recording schemes (Sharrock, 1976; Gibbons *et al*., 1993). Thus, we assumed that each surveyed grid square in which a species was not recorded represented an absence. However, preliminary analysis indicated that model fit was particularly low in coastal grid squares with very little land cover. Based on these results, we excluded grid squares with less than 10% land cover. We therefore proceeded to analyse presence-absence data for 112 bird species across 2603 of Great Britain’s 10–km grid squares at two time periods.

### Observed range changes

We compared species’ occupancy (*y*) between *t*_*1*_ and *t*_*2*_ across grid squares to identify observed changes in occupancy (*Δy*) – including instances of gain (where *y*_*t1*_ = 0 and *y*_*t2*_ = 1), persistence (where *y*_*t1*_ = 1 and *y*_*t2*_ = 1) and loss (where *y*_*t1*_ = 1 and *y*_*t2*_ = 0) – as well as areas that remained unoccupied (where *y*_*t1*_ = 0 and *y*_*t2*_ = 0).

### Climate predictors

We obtained data on four climate variables – mean temperature of the coldest month (°C), mean temperature of the warmest month (°C), ratio of actual to potential evapotranspiration (standard moisture index), and total annual precipitation (mm) – from the Climate Research Unit ts2.1 (Mitchell & Jones, 2005) and the Climate Research Unit 61-90 (New *et al*., 1999). We chose these variables to reflect known climatic constraints on bird distributions (Lennon *et al*., 2000; Illán *et al*., 2014). In each grid square, we calculated the mean value of each predictor over the periods 1966 – 1972 and 1986 – 1991, corresponding to *t*_*1*_ and *t*_*2*_, respectively, with two years tagged onto the start. We included these additional years since the presence-absence of birds in a particular breeding season is likely to depend on the climate of previous years (Araújo *et al*., 2005; Bradshaw *et al*., 2014).

### Climatic niches and climate redistribution

We estimated the realised climatic niches of bird species by correlating presence-absence data with climate variables in period *t*_*1*_ using generalised boosted models (GBMs; Ridgeway 1999). We chose GBMs as they were the most temporally-transferable single method in a previous study of climatic associations in British birds (Rapacciuolo *et al*., 2012) and perform consistently-well in additional studies of temporal transferability (Dobrowski *et al*., 2011; Smith *et al*., 2013). We fitted these models using the *gbm* package (Ridgeway, 2013) in R version 3.1.3 (R Core Team, 2014). We used custom code provided by Elith *et al*., (2008) to identify the optimal number of trees to be fitted in each model and avoid over-fitting calibration data. This code performs a 10-fold cross-validation procedure for each 50-tree increment, checking for improvements in calculated deviance on held-out data. Final models were fitted using the optimal number of trees identified through cross-validation (with a minimum of 1000 trees), 5 nodes, a learning rate of 0.001, and a bag fraction of 0.5. We assessed model fit in *t*_*1*_ using the area under curve (AUC) of the receiver operating characteristic function (Hanley & McNeil, 1982) – a measure of discrimination – and the point biserial correlation (COR) (Elith *et al*., 2006) – the Pearson correlation between observations and predictions. We calculated these measures of fit by averaging their values over each of the 10 folds held out during model calibration.

We used the realised climatic niches identified in *t*_*1*_ to generate (i) modelled estimates of probability of presence in *t*_1_ (*m*_*t1*_) based on climate predictor values for that period, and (ii) modelled estimates of probability of presence in *t*_2_ (*m*_*t2*_) after updating climate predictor values to reflect the redistribution of climatic conditions in *t*_*2*_. We then estimated change in modelled probability of presence given the redistribution of climatic conditions Δ(*m*) by subtracting *m*_*t1*_ from *m*_*t2*_. It is important to note that the predicted probability that a species will shift its range is not only conditional on its modelled change in probability of presence but also on its baseline probability of presence (Rapacciuolo *et al*., 2014b). As a result, we weighted Δ*m* values relative to *m*_*t1*_ (Δ*m*_*w*_; calculated by dividing negative Δ*m* values by *m*_*t1*_ and positive Δ*m* values by 1-*m*_*t1*_). Δ*m*_*w*_ values range from -1 – a 100% loss in predicted probability of presence to 1 – a 100% gain in probability of presence.

### Climatic niche tracking

#### Temporal validation plots

We estimated the relationship between observed changes in occupancy (Δ*y*) and predicted changes given climate redistribution (Δ*m*_*w*_) throughout the study area using temporal validation (TV) plots (Rapacciuolo *et al*., 2014b). The approach of TV plots is illustrated in Figure 1. For a given species, TV plots quantify the agreement between the probability of observing instances of loss, persistence, or gain (collectively, Δ*y* values) and changes in modelled probability of presence given the redistribution of climate variables (negative and positive Δ*m*_*w*_ values) throughout study sites. They do so by fitting two non-parametric functions with a logit link. The *loss* function (red line; Fig. 1c) models the probability that a grid square is lost from the species’ distribution (1; red tick marks in bottom rug plot of Fig. 1c) or not (0; all non-loss observations, expect stable absences, which cannot experience additional loss) as a function of Δ*m*_*w*_ values. In parallel, the *gain* function (blue line; Fig. 1c) models the probability that a grid square has been gained (1; blue tick marks in top rug plot of Fig. 1c) or not (0; all non-gain observations, expect persistence observations, which cannot experience additional gain) as a function of Δ*m*_*w*_ values. By subtracting the loss from the gain function to calculate a single curve (continuous black line; Fig. 1c), TV plots estimate the relative probability that sites are observed to be gained, remain stable (neither gained nor lost), or be lost for any given value of Δ*m*_*w*_ across the modelled range of Δ*m*_*w*_ values (see Rapacciuolo *et al*., 2014b for additional details).

**Figure 1.**
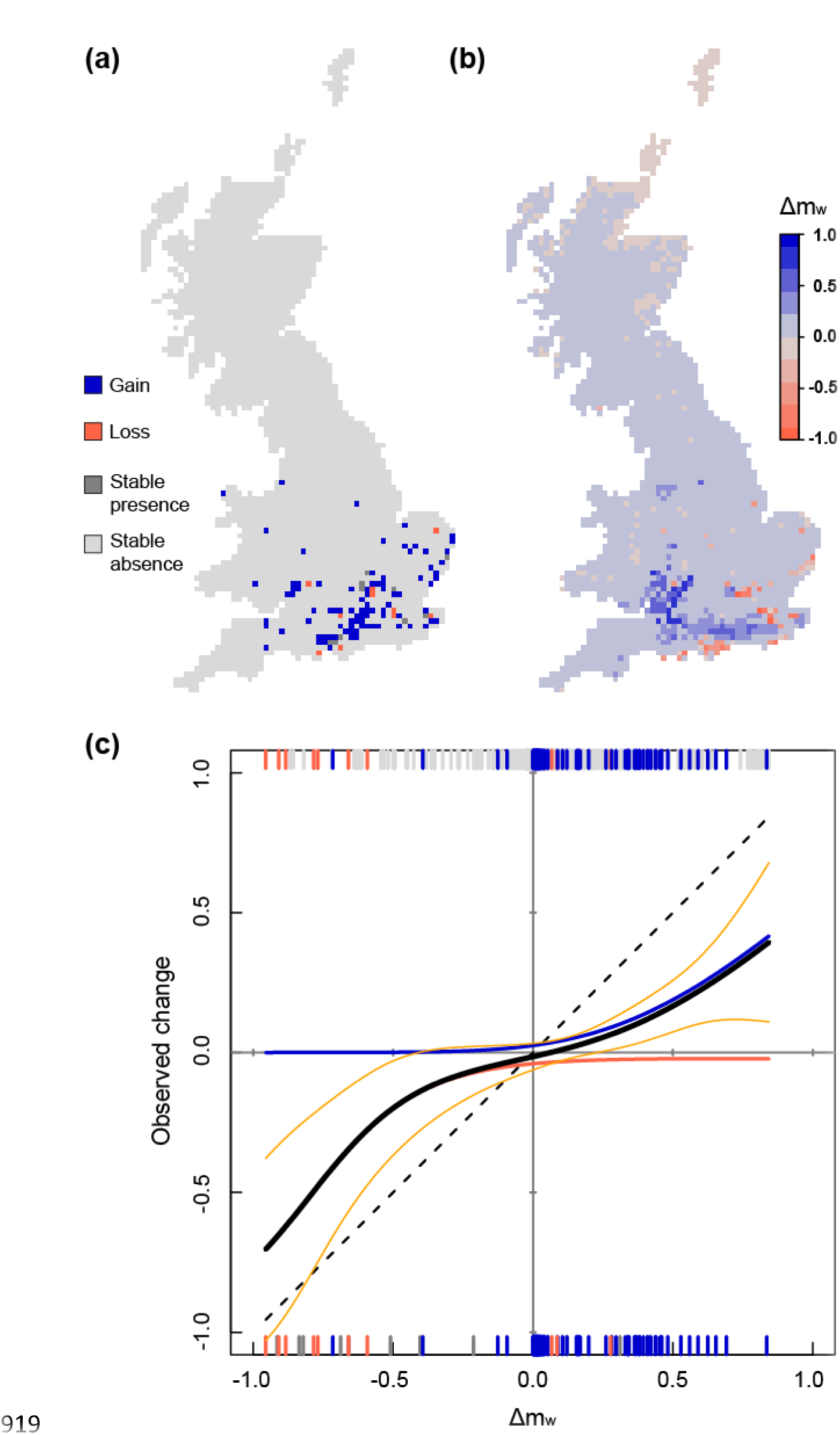
The approach of temporal validation (TV) plots exemplified using observations and model predictions for the Firecrest (*Regulus ignicapillus*). (a) Observed changes in the distribution of the Firecrest between *t*_*1*_ and *t*_*2*_, including observed gains (blue), losses (red), stable presences (dark grey), and stable absences (light grey). (b) Weighted changes in modelled probability of presence (Δ*m*_*w*_) for the Firecrest between *t*_*1*_ and *t*_*2*_. Δ*m*_*w*_values are derived by projecting in *t*_*2*_a model calibrated using presence-absence and climate data in *t*_*1*_. Bluer and redder colours indicate increases and decreases in probability of presence, respectively. (c) TV plot of the agreement between Δ*m*_*w*_ values from the climate-based SDM and observed changes for the Firecrest. Shown are the model temporal validation curve (thick black) – the sum of the plotted gain function (blue curve) and loss function (red curve) – and confidence intervals of ± 2 standard errors of the mean (orange). The dashed black line represents the ideal expectation for a perfect temporal validation curve. The rug plots show model values at observed sites; colours shown correspond to colours in panel (a).

#### Measuring climatic niche tracking

Assuming that changes in climate fully drive observed range changes and the processes of local gain and loss are unlimited and instantaneous (i.e. there are no time lags) every site with a predicted Δ*m*_*w*_ value of -1 should be observed to be lost whilst every site with a predicted Δ*m*_*w*_ value of 1 should be observed to be gained. Although there is an infinite number of monotonically-increasing curves connecting these two points, an ideal expectation for perfect niche tracking can be defined as a 1:1 line between observed and predicted changes passing from the origin (dashed black line; Fig. 1c). This line represents an ideal expectation for perfect niche tracking since it reflects the condition where every modelled Δ*m*_*w*_ value exactly equals the probability of observing a given change.

Based on this assumption, we quantified climatic niche tracking using Rapacciuolo *et al*. (2014b)’s accuracy of temporal validation (Acc_TV_), which accounts for the deviation between the ideal expectation and the modelled relationship between observed and predicted changes (the TV curve). Acc_TV_ is given by the mean absolute deviation between the ideal and the TV curve across all grid squares (Fig. 2), subtracted from 1 (Rapacciuolo *et al*., 2014b). Acc_TV_ values of 1 indicate perfect climatic niche tracking, whilst values < 1 indicate progressively lower tracking.

**Figure 2.**
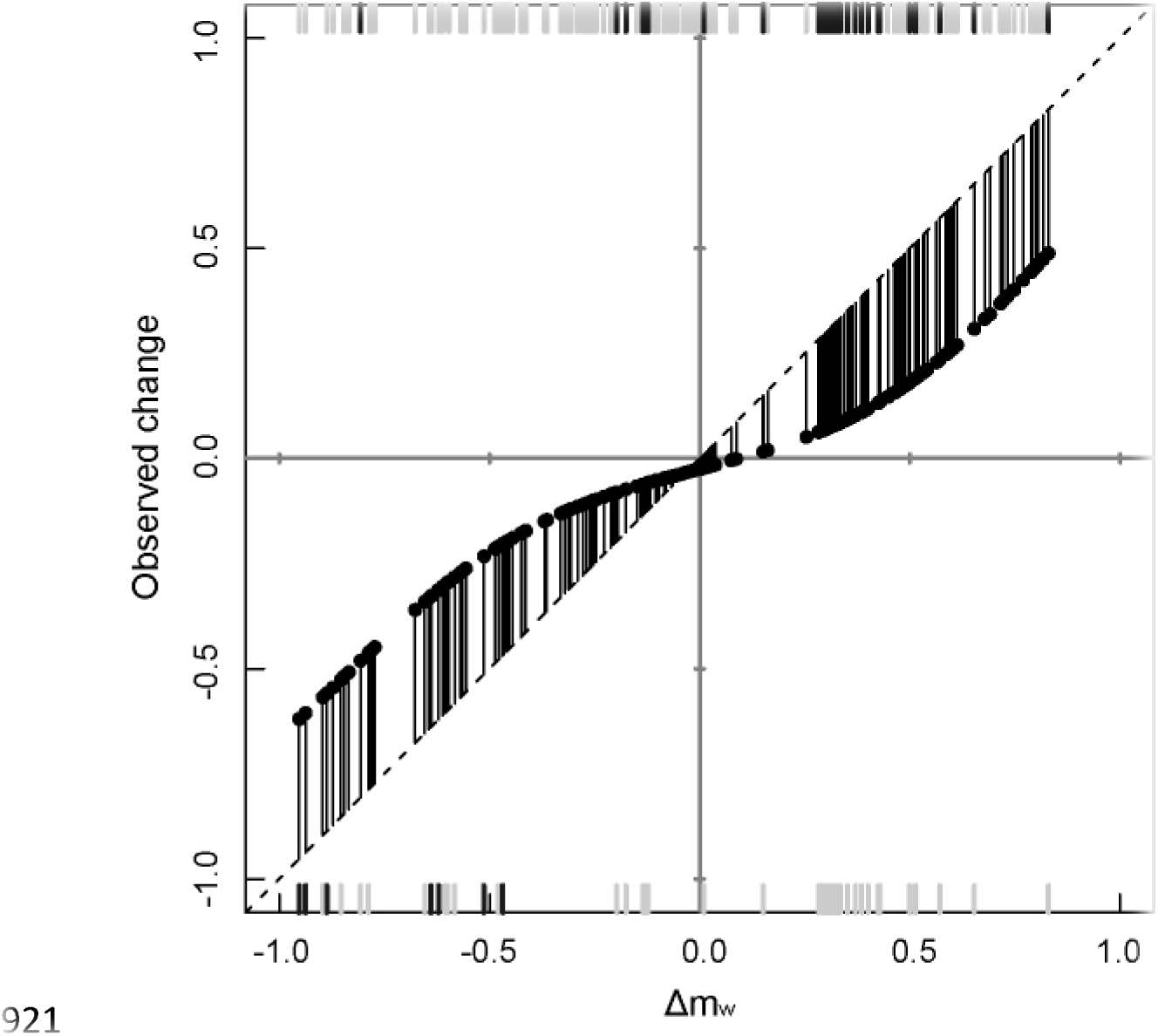
Measuring climatic niche tracking using temporal validation accuracy (Acc_TV_). Shown is a plot of observed range changes as a function of weighted changes in modelled probability of presence (Δ*m*_*w*_) for the Firecrest (analogous to Fig. 1c). Acc_TV_ is the mean absolute distance between the modelled y values (points) and the ideal *y* values (dashed black line), weighted by the corresponding absolute Δ*m*_*w*_ values at each observed site (tick marks), subtracted from 1. Data points from the Firecrest model were rarefied for ease of visualisation.

We tested whether Acc_TV_ values derived from temporal validation plots reliably measured climatic niche tracking using simulation (Appendix S1). We simulated range changes in a virtual species over a 2600-site artificial landscape based on change in two uniformly-distributed random climate covariates. We simulated varying scenarios of climatic niche tracking by modifying the degree to which range changes in the virtual species were determined by the specified functional response to climate. As expected, when the specified climate functional response fully determined the virtual species’ range changes (i.e. perfect climate niche tracking), Acc_TV_ values had a mean (± standard deviation) of 0.94 ± 0.01 (based on 999 simulation runs; Appendix S1, Fig. 1). Acc_TV_ values decreased progressively with climatic niche tracking; values of 0.41 ± 0.03 were associated with scenarios where 100% of the virtual species’ range changes were random with respect to climate change.

Since temporal validation plots use changes in modelled probability of presence weighted by baseline probability of presence (*m*_*t1*_), they may be sensitive to errors in model calibration in *t*_*1*_. For instance, say we have a site where *m*_*t1*_ = 0.8 but the species is absent in *t*_*1*_ (*y*_*t1*_ = 0): even a small increase in probability of presence in *t*_*2*_ (*Δm* = 0.1) will lead to a large weighted modelled change (*Δm_w_* = 0.1/(1 − 0.8) = 0.5) and, thus, a large deviation from observed change if the species remains absent (*y*_*t2*_ = 0). As a result, we also used our simulation to examine the effect of calibration errors on Acc_TV_ values (Appendix S1). Keeping the degree of niche tracking constant, we found that Acc_TV_ values were indeed sensitive to calibration errors and decreased with calibration accuracy (Appendix S1, Fig. S2). However, relatively large errors in model calibration (AUC = 0.70 ± 0.01; COR = 0.36 ± 0.02) were necessary to substantially affect Acc_TV_ values (≤ 0.85) when tracking was perfect. Thus, to remove the confounding effect of calibration error on Acc_TV_ values, we selected conservative thresholds for *t*_*1*_ AUC and COR representing acceptable calibration errors based on our simulations (AUC = 0.8; COR = 0.4). We then excluded all species with calibration AUC and COR values below these thresholds (18 out of 112 species).

### Effect of phylogeny and traits on climatic niche tracking

#### Phylogenetic signal

We used a recently-published molecular phylogeny (Thomas, 2008; Cassey *et al*., 2012) to identify evolutionary relationships among 109 species from the full set of 112. We tested whether closely-related species tended to have more similar Acc_TV_ values than species drawn at random from the phylogeny by estimating the maximum likelihood value of Pagel’s λ (Pagel, 1999). λ measures the agreement between observed trait variation across a phylogeny and a pure Brownian model of evolution (Freckleton *et al*., 2002); it ranges from 0 for phylogenetic independence to 1 for phylogenetic dependence. Importantly, we accounted for measurement error in Acc_TV_ values by incorporating within-species standard errors in our estimation of λ (Ives *et al*., 2007). We estimated λ values using the function phylosig in the R package phytools (Revell, 2012).

#### Biological traits

To test our four trait-based hypotheses, we obtained data on four biological traits of British birds: natal dispersal, adult survival, trophic level and species specialization index (SSI). We obtained natal dispersal estimates (in km) from Barbet-Massin *et al*. (2012). These estimates were obtained directly or extrapolated from published estimates of mean straight-line distance (in km) between the location birds were ringed in their year of birth and the location in which they were recovered at first breeding age (Paradis *et al*., 1998). We chose adult survival – calculated as the average proportion of birds of breeding age surviving each year (Robinson 2005) – as our measure of life-history speed. We also considered body size and reproductive output as additional measures of life-history speed but, given the high inter-correlation among the three variables, we only kept adult survival. We generated a factor variable for trophic level by placing each species into one of 5 categories (modified from Huntley *et al*. 2004): (i) exclusively herbivorous species; (ii) herbivorous/insectivorous species, with predominantly herbivorous diet; (iii) herbivorous/insectivorous species, with predominantly insectivorous diet; (iv) insectivorous species and carnivorous species predominantly consuming herbivorous prey; (v) carnivorous species predominantly consuming carnivorous prey. Finally, we estimated species’ habitat specialization using the species specialization index (SSI), a measure of evenness in habitat affinity (Devictor *et al*., 2008b). The higher the SSI, the more specialised a species. SSI values were calculated by Le Viol *et al*. (2012) for 99 of the species in our final dataset, based on the coefficient of variation in habitat affinity across 98 habitat categories in Europe (Le Viol *et al*., 2012).

#### Trait models

We examined whether biological traits could predict variation in climatic niche tracking, as measured by Acc_TV_. Because shared natural history among our set of species unaccounted by the modelled traits may lead more phylogenetically-related species to respond more similarly, modelling individual species as statistically-independent units may lead to biased results. Therefore, we accounted for shared phylogenetic history in our trait models using phylogenetic generalised least squares (PGLS) models – as implemented in the R package CAPER (Orme *et al*., 2011) – which incorporate covariances between species into the model’s error term. To avoid under-or over-correcting for phylogenetic autocorrelation, we estimated the degree of phylogenetic dependence in model residuals by estimating the maximum-likelihood value of Pagel’s λ (Pagel, 1999) simultaneously with the other model parameters.

We constructed a PGLS model set including all possible combinations of the single and additive effects of natal dispersal, adult survival, trophic level and SSI, as well as an intercept-only model. We standardised all continuous predictors in each model (by subtracting the mean and dividing by the standard deviation); effect sizes obtained this way provide a measure of the importance of each predictor on the response (Schielzeth, 2010). All PGLS models assumed normally-distributed model residuals; visual inspection of residuals vs fitted values plots and quantile-quantile plots confirmed that no model violated this assumption.

In order to derive reliable estimates of the sign and magnitude of the effect of each predictor based on the full set of potential trait models, we employed multimodel inference (Burnham & Anderson, 2004; Johnson & Omland, 2004). We first ranked all potential models using the Akaike Information Criterion correction for small sample sizes (AICc; Burnham and Anderson 2002). For each model in the full set, we quantified the probability that it was the best model given the data using AICc weights (AIC_w_), and its structural goodness-of-fit using adjusted R^2^. Taking each predictor in turn, we then considered the full set of models in which the predictor appeared and calculated: i) its relative importance, by summing the AIC_w_ values across the model set (∑AIC_w_) and ii) model-averaged coefficients and standard errors by averaging coefficients across all models in the set that included the focal variable, weighted by each model’s AIC_w_ (Johnson & Omland, 2004). For predictor coefficient averages, AIC_w_ values were recalculated over all models in which each predictor appeared, in order to make sure AIC_w_ values used for weighting added up to 1.

## RESULTS

### Climatic niches and climate redistribution

When assessed against held out presence-absence data in *t*_1_, our models showed excellent discrimination (AUC; mean ± standard deviation = 0.90 ± 0.06; see Fig. S1 in supporting information) and correlation (COR; 0.60 ± 0.20). However, 18 (out of 112) species did exceed our simulation-based thresholds for acceptable error during model calibration (AUC < 0.8; COR < 0.4), so we only considered the remaining 94 species in further analyses.

When projected on updated climate values in *t*_2_, the mean discriminatory power and correlation of our models both decreased (AUC: 0.86 ± 0.08; COR: 0.53 ± 0.17; Fig. S1). We examined the pattern of grid square-wise mean predicted change in probability of presence (Δ*m*_*w*_) across all species and found that the majority of grid squares across Great Britain were predicted to have a positive mean Δ*m*_*w*_ (i.e. overall gains; see Fig. S2). Mean Δ*m*_*w*_ values were highest in the highlands of Wales and western Scotland – where total precipitation increased most and standard moisture decreased least (Figure S3a, b) – and lowest in the Shetland Islands and south-eastern England – where mean temperatures increased most (Fig. S3c, d).

#### Climatic niche tracking

The degree of climatic niche tracking among the 94 British bird species was low overall (Acc_TV_: 0.52 ± 0.20; Fig. 3). When compared with our simulation results, the observed mean Acc_TV_ for British birds approached the value derived from scenarios where only 10% of the virtual species’ range changes were determined by climate (Appendix S1, Fig. S1). However, observed Acc_TV_ values varied considerably among bird species, with a number of species tracking their climatic niches closely and others shifting their ranges irrespective of or even opposite to climatic expectations (Fig. 3).

**Figure 3.**
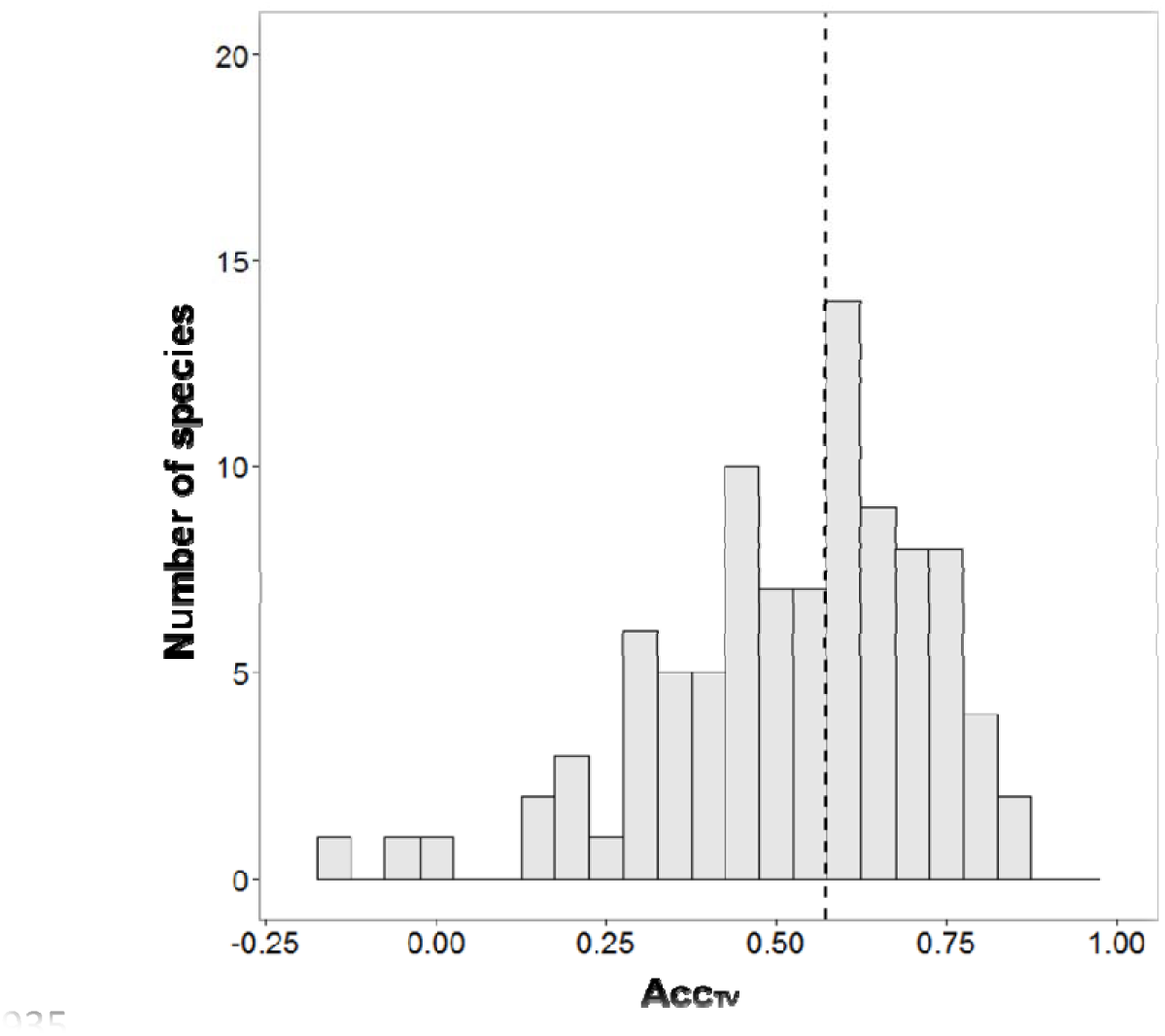
Distribution of Acc_TV_values across 94 species of British breeding birds. Acc_TV_ is a measure of climatic niche tracking; values of 1 indicate perfect niche tracking. The dashed line indicates the median Acc_TV_ across all species (0.583). Acc_TV_ values for species with high errors during model calibration were excluded from this analyses (see Methods section).

#### Effect of phylogeny and traits on climatic niche tracking

We limited our comparative analyses to 70 species with complete phylogenetic and trait information, as well as low calibration error (there was no significant difference in Acc_TV_ distribution between this subset and the set of 94 species of Fig. 3; *t*_144_ = −0.54, *p* = 0.59; Fig. S4). After accounting for uncertainty due to measurement error, the phylogenetic signal in Acc_TV_ values was not significantly different from 0 (λ = 0; p = 1). Although a low phylogenetic signal may suggest the use of PGLS models is unwarranted, the phylogenetic signal in the residuals of trait models was not null (upper 95% confidence intervals of maximum-likelihood lambda values across trait models ranged from 0.17 – 0.29; Table 1). As a result, we proceeded by running phylogenetic trait models and present the results from these models below. However, we also ran non-phylogenetic generalised linear models (GLMs) for comparison. Given the minimal phylogenetic correction applied in PGLS models (Table 1), differences from GLMs were negligible (Tables S2, S3).

**Table 1.** Summary of model selection for phylogenetic generalised least squares(PGLS) models of climatic niche tracking (Acc_TV_) as a function of biological traits in British birds. Traits considered were adult survival (Surv), trophic level (Troph), natal dispersal (Disp), and habitat specialization (species specialization index; SSI).

| Acc<sub>TV</sub> PGLS models |  |  |  |  |  |  |  |  |  |  |  |  |  |  |  |  |
| --- | --- | --- | --- | --- | --- | --- | --- | --- | --- | --- | --- | --- | --- | --- | --- | --- |
|  | Model rank |  |  |  |  |  |  |  |  |  |  |  |  |  |  |  |
|  | 1 | 2 | 3 | 4 | 5 | 6 | 7 | 8 | 9 | 10 | 11 | 12 | 13 | 14 | 15 | 16 |
| Surv | • | • | • | • | • | • | • | • |  |  |  |  |  |  |  |  |
| Troph | • | • |  |  | • |  |  | • |  |  |  | • |  | • | • | • |
| Disp |  | • | • |  |  | • |  | • |  | • |  |  | • | • |  | • |
| Spec |  |  |  |  | • | • | • | • |  |  | • |  | • |  | • | • |
| **ΔAIC** | 0.0 | 1.4 | 1.5 | 1.6 | 2.4 | 3.3 | 3.6 | 3.7 | 4.3 | 5.6 | 6.4 | 6.5 | 7.7 | 8.7 | 8.9 | 11.2 |
| **LL** | 17.4 | 17.9 | 13.1 | 12.0 | 17.4 | 13.3 | 12.1 | 18.0 | 9.6 | 10.0 | 9.6 | 12.9 | 10.0 | 13.0 | 12.9 | 13.0 |
| **AIC<sub>w</sub>** | 0.28 | 0.14 | 0.13 | 0.13 | 0.09 | 0.05 | 0.05 | 0.05 | 0.03 | 0.02 | 0.01 | 0.01 | 0.00 | 0.00 | 0.00 | 0.00 |
| **λ<sub>upper</sub>** | 0.12 | 0.12 | 0.17 | 0.15 | 0.12 | 0.17 | 0.15 | 0.13 | 0.20 | 0.28 | 0.21 | 0.20 | 0.29 | 0.24 | 0.20 | 0.25 |
| **R<sup>2</sup>** | 0.14 | 0.14 | 0.07 | 0.05 | 0.12 | 0.06 | 0.04 | 0.13 | 0.00 | 0.00 | 0.00 | 0.04 | 0.00 | 0.02 | 0.02 | 0.01 |

*Notes:* the variables included in each model are shown with the symbol •. Models are ranked in order of increasing AICc differences (ΔAIC). The log likelihood (LL) and Akaike weights (AIC<sub>w</sub>) indicate the relative likelihood of a model given the data, λ<sub>upper</sub> represents the 95% upper confidence interval for the maximum-likelihood value of phylogenetic dependence in the model residuals (all maximum-likelihood λ means were 0), and R<sup>2</sup> indicates the proportion of the total variation in Acc<sub>TV</sub> explained by the model predictors. All models were built using 70 species with reliable climatic niche tracking measures complete phylogenetic and trait information.

The best-supported trait model had a relatively low AICc weight (AIC_w_ = 0.283; Tables 1, S2), indicating there was no overwhelming support towards any particular trait model (Johnson & Omland, 2004). Overall, models incorporating phylogeny and traits explained a very small portion of variation in Acc_TV_ values, up to a maximum adjusted R^2^ of 0.14 (mean-adjusted R^2^ ± standard deviation: 0.050 ± 0.054; Tables 1, S2).

Relative importance values supported adult survival as the most important trait predictor of Acc_TV_ (∑AIC_w_ = 0.91; Table 2), with model-averaged coefficients indicating a negative effect of adult survival on Acc_TV_ (Table 2). Furthermore, trophic level was also an important predictor of AccTV (∑AIC_w_ = 0.57) species with a mixed herbivorous/insectivorous diet had lower AccTV values compared to exclusively-herbivorous and exclusively-carnivorous species (Table 2). We found no support for an effect of natal dispersal or SSI on Acc_TV_ (Tables 2, S4).

**Table 2.** Summed AIC weight (∑AIC_w_) and model-averaged coefficient for each trait predictor of climatic niche tracking (Acc_TV_) in British birds across the full set of phylogenetic generalised least squares (PGLS) models. The standard error for each coefficient estimate is indicated in brackets. Average coefficients with confidence limits not overlapping zero are shown in boldface.

| $Acc_{TV}$ PGLS models | | |
| --- | --- | --- |
| | $\sum AIC_w$ | Model-averaged coefficient |
| (Intercept) | – | **0.722 (0.252, 1.192)** |
| Survival | 0.91 | **-0.069 (-0.121, -0.017)** |
| Trophic level | 0.57 | – |
| 2 | – | **-0.473 (-0.882, -0.064)** |
| 3 | – | **-0.463 (-0.920, -0.006)** |
| 4 | – | -0.333 (-0.736, 0.070) |
| 5 | – | -0.327 (-0.783, 0.130) |
| Dispersal | 0.40 | 0.031 (-0.021, 0.083) |
| Specialization | 0.25 | -0.012 (-0.074, 0.049) |

*Notes:* Trophic levels are coded as follows: 2 = herbivorous/insectivorous species with predominantly-herbivorous diet, 3 = herbivorous/insectivorous species with predominantly-insectivorous diet, 4 = insectivorous species and carnivorous species mostly consuming herbivorous prey; 5 = carnivorous species mostly consuming carnivorous prey. All models were built using 70 species with reliable climatic niche tracking measures and complete phylogenetic and trait information.

## DISCUSSION

Evidence that species are responding individualistically to the same changes in climate (Chen *et al*., 2011; Rapacciuolo *et al*., 2014a) highlights the key role that biological traits play in determining distributional responses to climate change (Williams *et al*., 2008; O’Connor *et al*., 2012; Foden *et al*., 2013). By comparing the redistribution of species’ climatic associations with their recently-observed range gains and losses, we were able to test a number of hypotheses of the effect of biological traits on species’ climatic niche tracking.

Overall, our results indicate that British breeding birds did not track their climatic niches closely and observed species’ range shifts deviated substantially from climate change expectations over an approximately 30-year period. However, there was high heterogeneity among species in their degree of climatic niche tracking. A number of species, whose demographic rates are known to be significantly impacted by climate, did show a relatively high degree of climatic niche tracking. These included the Pied White Wagtail (*Motacilla alba*), whose first egg dates and juvenile survival rates increase with spring temperatures (Mason & Lyczynski, 1980; Crick & Sparks, 1999), the Merlin (*Falco columbarius*), whose regional declines have previously been linked with climate change drivers (Ewing *et al*. 2011), and the Blackcap (*Sylvia atricapilla*), whose overwinter survival rates have been improved by milder winter conditions (Plummer *et al*. 2015). In contrast, several other species appeared to have shiftedi rrespective of, or even counter to, climate redistribution. Previous studies over similar timescales also found high heterogeneity in the degree of climatic niche tracking across bird species (Gregory *et al*., 2005; Green *et al*., 2008; Maggini *et al*., 2011; La Sorte & Jetz, 2012). One possible explanation for this pattern is that some species’ distributional responses may lag behind climate change (Menéndez *et al*., 2006; Devictor *et al*., 2008a). Indeed, studies over longer timescales suggest that, given enough time, the overall degree of climatic niche tracking is generally higher (e.g. Tingley *et al*. 2009, 2012). Alternatively, observed distribution changes of British breeding birds over our study period may not have been primarily driven by climate. For instance, population declines and range contractions in a number of British bird species are thought to be a consequence of changes in land-use (Thomas *et al*., 2004; Eglington & Pearce-Higgins, 2012). This explains why species such as the Nightingale and the Turtle Dove – which have been hugely impacted by agricultural intensification and changing farming practices (Fuller *et al*., 1995; Browne *et al*., 2004) – displayed the lowest degree of climatic niche tracking. Lag effects and alternative drivers of change are only two of the potential explanations for mismatches between observations and climate-based predictions. Those and additional factors – such as changing biotic interactions – are undoubtedly required for a full attribution of observed range shifts. However, a full attribution of the drivers of recent range shifts was beyond the scope of our study, which instead focused on distinguishing species whose changes were consistent with climate predictions from species requiring additional processes. With this objective in mind, we believe that temporal validation plots and associated measures such as Acc_TV_ are a useful tool and that their utility should increase as more temporal datasets of species’ distribution shifts become available.

Models incorporating both species’ traits and phylogeny explained only a small portion of the variation in climatic niche tracking among British breeding birds. This is in line with previous studies of the effect of traits on measures of the agreement between climate-based predictions and observations (McPherson & Jetz, 2007; Angert *et al*., 2011; Smith *et al*., 2013). In general, species’ responses to climate change are likely to be complex, idiosyncratic and difficult to predict given the multitude of interacting biological and environmental factors underlying them (Pimm 2009; Walther 2010; LaSorte and Jetz 2012). Our models were over-simplistic – limited to a number of hypotheses based on solid theoretical foundations – and should undoubtedly include additional processes. For instance, behavioural attributes such as activity times and nesting behaviour have been posited as important predictors of variation in climate change responses in mammals (McCain & King, 2014) and represent a fruitful direction for further theoretical and empirical work. Furthermore, an approach that directly tests the effects of species’ biological traits on climatic niche tracking may be preferable or at least complementary with the indirect statistic on statistic approach we use here. However, it is not obvious how one would develop such direct approach without incurring a significant loss of information from the calculation of assemblage-level trait summaries (e.g. Douma *et al*., 2012).

Together with the general challenges shared among studies of climate change responses, a number of factors specific to our study system may underlie the low explanatory power of our models. Although the British breeding bird data we use here are among the highest quality datasets on spatiotemporal biodiversity changes, their temporal and spatial extents may not be sufficient to detect climatic niche tracking. First, a 30-year time interval may not be sufficient to detect substantial distributional responses to climate changes for most British breeding bird species. While this may be due to the aforementioned lag effects, it may also simply result from the fact that climatic conditions in Britain may not have changed sufficiently to generate a response for most species. Acc_TV_ estimates may be particularly prone to error for species experiencing lower magnitudes and extents of climate change. For instance, lower magnitudes and extents of climate change have been found to bias Acc_TV_ towards higher values by leading to intrinsically-lower mean deviations between predictions and observations (Rapacciuolo *et al*., 2014b). Despite the low correlation of Acc_TV_ with both magnitude (measured as the range of Δ*m*_*w*_ values; ρ = 0.12) and extent of change (measured as the total number of observed gains and losses; ρ = 0.10), we acknowledge that variation in these species-specific aspects of climate change exposure may still have impacted Acc_TV_ values. In general, we do caution against the use of temporal validation plots and Acc_TV_ for comparing among species and geographical areas with radically different climate change exposures. A second shortcoming of our particular study system is that Britain may not be a sufficient spatial extent to detect climate change responses for the species in our dataset, all of which have breeding ranges extending beyond Britain. Furthermore, Britain constitutes the northwestern boundary for many of these species’ ranges and may not accurately reflect the entire spectrum of climatic conditions they can occupy. An important consequence of this is that the climatic niches we estimated are likely to be incomplete for some species. We acknowledge that the failure to capture the full extent of species’ climatic niches may be partially responsible for the deviations we identified between observed and predicted distribution changes. However, we preferred limiting our study to the standardised British data rather than incorporating additional European data on the species’ ranges (e.g. EBCC Atlas of European Breeding Birds; Hagemeijer & Blair, 1997) to avoid the perils of integrating data across different spatial and temporal scales (McPherson *et al*., 2006; Bombi & D’Amen, 2012). These factors considered, the British breeding bird dataset we used here may appear as an unsuitable choice for testing hypotheses of the effect of traits on climatic niche tracking. However, it is one the few and, arguably, one of the highest-quality datasets that enables performing such tests. If hypotheses of climatic niche tracking are not testable using the best datasets currently available, they are in danger of not being testable at this time.

Our models did provide evidence that life-history speed and trophic level were the most important predictors of climatic niche tracking we considered. As hypothesised, species with lower adult survival were more likely to have tracked their climatic niches over the time period of our study. A likely explanation for this is that short generation times and higher rates of population growth lead to a higher likelihood of rapid expansion and subsequent establishment into newly-suitable areas (Angert *et al*., 2011; Anderson *et al*., 2012; O’Connor *et al*., 2012; Schloss *et al*., 2012). Our result is in line with recent findings that life-history speed is positively correlated with population increase (Robinson *et al*., 2014) and range expansion (Bradshaw *et al*., 2014) in British birds. Conversely, our hypothesis that increasing trophic level would lead to lower climatic niche tracking due to increasing separation from direct climatic effects was only partially supported. Species from both the lowest (i.e. exclusively-herbivorous species) and the highest (i.e. exclusively-insectivorous/carnivorous species) trophic levels tracked their climatic niches more closely than species from intermediate trophic levels (i.e. mixed herbivorous/insectivorous species). In addition to our original hypothesis, a number of processes may underlie this result. For instance, evidence from mammals suggests that carnivores may be better able to track their climatic niches than herbivores and omnivores due to their higher dispersal velocity (Schloss *et al*., 2012) and wider range areas (Carbone *et al*. 2005). Furthermore, our measure of trophic level may have partially captured species’ differences in ecological generalisation, with mixed-diet generalists potentially displaying a lower degree of climatic niche tracking due to their lower susceptibility to climate change (Foden *et al*., 2013). Therefore, although we did not find evidence of an effect of natal dispersal or habitat specialisation on climatic niche tracking, it is possible that trophic level may have indirectly captured part of their hypothesised effects.

A further noteworthy result was that the phylogenetic signal in climatic niche tracking was not significantly different from zero, suggesting that biogeographic responses to climate change may be highly idiosyncratic among closely-related species. This pattern does not appear to be limited to British birds. A number of studies highlighted how congeneric species of birds and mammals are shifting their ranges in opposite directions (Moritz *et al*., 2008; Tingley *et al*., 2012; Rapacciuolo *et al*., 2014a). Moreover, several studies reported that accounting for phylogenetic relatedness among species did not modify their conclusions on the effect of traits on the performance of climate-based species distribution models (Green *et al*., 2008; Pöyry *et al*., 2008; Newbold *et al*., 2009). However, one study did find a weak but significant phylogenetic signal to the predicted suitable future climate of European species (Thuiller *et al*., 2011), which suggests that phylogeny remains an important factor to consider when assessing species’ vulnerability to climate change. At first glance, our finding of an extremely low phylogenetic signal appears at odds with the conclusions of Bradshaw *et al*., 2014, who found a mid-range phylogenetic signal in the change in area of occupancy for 106 British bird species (approximately 62 of which were shared with our 70-species subset; Bradshaw *et al*., 2014). However, our measure of climatic niche tracking Acc_TV_ was only weakly correlated with change in area of occupancy (ρ = 0.13), as it was based on local rather than whole-range area changes. As a result, there is no real reason to expect congruence in phylogenetic signal among these two studies.

Focusing on distribution changes consistent with climate change at the local scale can unveil patterns of species’ sensitivity to climate change which may not be identified by examining range changes as a whole. We present here a promising approach for doing so, which uses temporal validation plots and time series of distribution data to assess how well climate-based models predict observed distribution gains and losses at individual sites. Though we are unable to provide strong empirical evidence that biological traits mediate climatic niche tracking in this study, we believe our approach may prove to be useful in this context as biodiversity datasets at broad temporal and spatial extents become increasingly available.

## DATA ACCESSIBILITY

The species distribution data used in these analyses can be accessed via the National Biodiversity Network Gateway (1968–1972: https://data.nbn.org.uk/Datasets/GA000600; 1988–1991: https://data.nbn.org.uk/Datasets/GA000147). The climate data can be accessed via the Climate Research Unit (http://www.cru.uea.ac.uk/cru/data/hrg/). The bird phylogeny can be accessed from the relevant publications (Thomas, 2008; Cassey *et al*., 2012). R code to generate temporal validation plots can be found at https://github.com/giorap/tv-plots.

## ACKNOWLEDGEMENTS

We thank all volunteers contributing the atlas records on which this study is based. We thank David B. Roy and Albert B. Phillimore for useful comments on previous versions of this manuscript. GR received funding from the W.M. Keck Foundation and the Biotechnology and Biological Sciences Research Council (BBSRC). This paper is a contribution from the Imperial College Grand Challenges in Ecosystems and the Environment Initiative and the Berkeley Initiative in Global Change Biology.

